# Antibiotic resistance alters the ability of *Pseudomonas aeruginosa* to invade the respiratory microbiome

**DOI:** 10.1101/2023.11.14.567137

**Authors:** Selina Lindon, Sarah Shah, Danna R. Gifford, Maria A. Gomis Font, Divjot Kaur, Antonio Oliver, R. Craig MacLean, Rachel M. Wheatley

## Abstract

The emergence and spread of antibiotic resistance in bacterial pathogens is a global health threat. One important unanswered question is how antibiotic resistance influences the ability of a pathogen to invade the host-associated microbiome. Here we investigate how antibiotic resistance impacts the ability of the opportunistic bacterial pathogen *Pseudomonas aeruginosa* to invade the respiratory microbiome, by measuring the ability of *P. aeruginosa* spontaneous antibiotic resistant mutants to invade pre-established cultures of commensal respiratory microbes. We find that commensal respiratory microbes tend to inhibit the growth of *P. aeruginosa*, and antibiotic resistance is a double-edged sword that can either help or hinder the ability of *P. aeruginosa* to overcome this inhibition. The directionality of this help or hinderance depends on both *P. aeruginosa* genotype and respiratory microbe identity. Antibiotic resistance facilitates the invasion of *P. aeruginosa* into *Staphylococcus lugdunensis,* yet impairs invasion into *Rothia mucilaginosa* and *Staphylococcus epidermidis*. *Streptococcus* species provide the strongest inhibition to *P. aeruginosa* invasion, and this is maintained regardless of antibiotic resistance genotype. Our study demonstrates how antibiotic resistance can alter the ability of a bacterial pathogen to invade the respiratory microbiome and suggests that attempts to manipulate the microbiome should focus on promoting the growth of commensals that can provide robust inhibition of both wildtype and antibiotic resistant pathogen strains.

## Introduction

Antibiotic resistance in pathogenic bacteria has emerged as a serious threat to public health (*1, 2*). Infections caused by antibiotic resistant pathogens are associated with worse clinical outcomes for patients, longer hospitalisations, and higher healthcare costs (*3*). Our microbiome can provide protection against colonisation by pathogenic bacteria. This microbiome-mediated colonisation resistance can be conferred through a variety of mechanisms, including induction of host immune responses, metabolic niche exclusion, and direct antagonistic interactions such as toxin production (*4*).

One key unresolved question is how antibiotic resistance impacts the ability of pathogens to colonise hosts. Antibiotic resistance is often associated with fitness costs (*5–7*), suggesting that resistant pathogens should have a decreased ability to invade host-associated microbiomes. Yet, many antibiotics were originally isolated from microbes or are modifications of microbial products and mediate inter-bacterial competition in their natural capacity (*8*). As such, mechanisms of antibiotic resistance may also provide cross-resistance to anti-competitor toxins produced by commensal microbes in the microbiome. In these scenarios, resistant pathogens may have an increased ability to invade the microbiome. To fully comprehend the consequences of antibiotic resistance in bacterial pathogens, we need to understand how antibiotic resistance impacts the ability of a pathogen to invade the microbiome, which has clear significance for the potential of resistant strains to transmit between patients.

Here we set out to probe this question of how resistance alters the invasion ability of a pathogen, by using the opportunistic bacterial pathogen *Pseudomonas aeruginosa* and the respiratory microbiome as our model system. *P. aeruginosa* can cause a wide range of infections but is particularly problematic in the lungs. *P. aeruginosa* is a major causative pathogen in serious short-term infections such as ventilator-associated pneumonia and is able to colonise the lungs of patients with bronchiectasis or cystic fibrosis, resulting in long-term recurring infections. In addition to widespread multidrug resistance (*9*), *P. aeruginosa* is also known for its impressive arsenal of virulence factors and anti-competitor weaponry (*10*). All of this helps *P. aeruginosa* function as a powerful invasive species that can cause difficult-to treat-infections associated with a high burden of mortality (*1*). The respiratory microbiome can provide protection against pathogen colonisation (reviewed recently here: (*11–15*)). In healthy individuals, the lung microbiota largely resembles the upper respiratory tract microbiota, but at lower densities (*16, 17*), and common constituent members include *Streptococcus*, *Staphylococcus*, *Prevotella*, *Veillonella* and *Rothia* species (*12, 18*).

In this study, we generated spontaneous resistant mutants of *P. aeruginosa* to three clinically important anti-pseudomonal drugs (ciprofloxacin, ceftazidime and meropenem). Both meropenem and ceftazidime target cell wall synthesis, whereas ciprofloxacin targets nucleic acid synthesis via DNA gyrase (*19*). We tested the ability of these mutants to invade pre-established cultures of six commensal respiratory microbiome strains. While these respiratory microbes tended to inhibit *P. aeruginosa* growth, we found that antibiotic resistance could either help or hinder the ability of *P. aeruginosa* to invade, and the directionality of this was dependant on both *P. aeruginosa* genotype and respiratory microbe identity.

## Methods

### Respiratory microbiome strains

The six respiratory microbiome species used were: *Staphylococcus epidermidis*, *Rothia mucilaginosa*, *Streptococcus gordonii*, *Staphylococcus lugdunensis*, *Streptococcus oralis*, and *Streptococcus salivarius* (Supplementary Table 1). These microbiome species were selected due to presence and relevance in the respiratory microbiome, availability of strain, and ability of strain to grow in proposed assay conditions.

### Generation of antibiotic resistant *P. aeruginosa* strains

A green fluorescent protein (GFP)-tagged strain of *P. aeruginosa* PAO1 (PAO1-GFP; (*20*)) was used so that reads in the GFP-channel could measure *P. aeruginosa* invasion (excitation: 485/20, emission: 516/20). This strain was previously created by insertion of GFP and a gentamicin resistance cassette into the Tn7 transposon insertion site of the reference strain of *P. aeruginosa* PAO1 (*20*). Three independent spontaneous resistant mutants were selected on each of the three antibiotics (ciprofloxacin, ceftazidime, and meropenem) at the clinical breakpoints according to the EUCAST Clinical Breakpoint Tables v.12.0 (ciprofloxacin 0.5 μg/mL, ceftazidime 8 μg/mL and meropenem 8 μg/mL) (*21*). These antibiotics were chosen to represent different classes of clinically important antipseudomonal drugs (fluoroquinolones, cephalosporins and carbapenems).

To generate spontaneous resistant mutants, PAO1-GFP was grown from glycerol stock on LB (Lysogeny Broth) Miller with agar (Sigma-Aldrich) plates supplemented with 15 μg/mL gentamicin overnight at 37 °C. Single colonies were grown in LB Miller broth overnight at 37 °C in biological triplicate with shaking at 225 rpm, then 150 uL of overnight culture was plated on LB Miller agar plates supplemented with either 0.5 μg/mL ciprofloxacin, 8 μg/mL ceftazidime, or 8 μg/mL meropenem. Spontaneous resistant mutants of PAO1-GFP successfully grew on both 0.5 μg/mL ciprofloxacin and 8 μg/mL ceftazidime. A single colony from each plate was inoculated into LB Miller broth supplemented with the corresponding antibiotic at 50% of the selective concentration (i.e. 0.25 μg/mL ciprofloxacin and 4 μg/mL ceftazidime) for overnight growth. Three resistant mutants for each antibiotic were stored as glycerol stocks at –80°C (Supplementary Table 1). No resistant mutants of PAO1-GFP were able to grow on meropenem 8 μg/mL, and so PAO1-GFP was serially passaged through progressively higher meropenem concentrations to generate meropenem resistant mutants, ramping from 1 μg/mL to 2 μg/mL to 4 μg/mL with transfer every 24 hours before plating on 8 μg/mL meropenem. The same procedure for stocking was then followed as above. A total of nine resistant mutants of PAO1-GFP were generated across the three antibiotics which are hereby referred to as: ciprofloxacin resistant mutant 1 (cipR1), ciprofloxacin resistant mutant 2 (cipR2), ciprofloxacin resistant mutant 3 (cipR3), ceftazidime resistant mutant 1 (cefR1), ceftazidime resistant mutant 2 (cefR2), ceftazidime resistant mutant 3 (cefR3), meropenem resistant mutant 1 (merR1), meropenem resistant mutant 2 (merR2) and meropenem resistant mutant 3 (merR3).

### Antibiotic resistance phenotyping via minimum inhibitory concentration assays

Isolates were grown from glycerol stocks on LB Miller Agar plates overnight at 37 °C. Single colonies were then inoculated into tryptic soy broth (Sigma-Aldrich) for overnight growth at 37 °C with shaking at 225 rpm, after which overnight suspensions were serial diluted to ∼5 × 10^5^ CFU/mL. Antibiotic resistance phenotyping was carried out as minimum inhibitory concentration (MIC) assays via broth microdilution as defined by EUCAST guidelines (*22*), with the alteration of tryptic soy broth for the growth media as this was the media used for the invasion assays. MIC was calculated using the following 2-fold dilution series for meropenem (0 μg/mL, 0.5 μg/mL, 1 μg/mL, 2 μg/mL, 4 μg/mL, 8 μg/mL, 16 μg/mL, 32 μg/mL, 64 μg/mL), ciprofloxacin (0 μg/mL, 0.03125 μg/mL, 0.0625 μg/mL, 0.125 μg/mL, 0.25 μg/mL, 0.5 μg/mL, 1 μg/mL, 2 μg/mL, 4 μg/mL), and ceftazidime (0 μg/mL, 0.5 μg/mL, 1 μg/mL, 2 μg/mL, 4 μg/mL, 8 μg/mL, 16 μg/mL, 32 μg/mL, 64 μg/mL) designed to capture ranges either side of the clinical breakpoint (*21*). Growth inhibition was defined as OD_595_ < 0.200 and we calculated the MIC of each isolate as the median MIC score from three biological independent assays of each isolate. For some strains, the MIC was the lowest concentration in this range. However, as we were using this assay to determine increased resistance around the clinical breakpoint that sat in the midpoint of these ranges, this was not an issue.

### Growth rate assays

Growth rate assays were used to characterise growth of the antibiotic resistant mutants compared to the wildtype (PAO1-GFP) in tryptic soy broth in the absence of any respiratory microbes. *P. aeruginosa* strains were grown from glycerol stock on LB Miller agar plates overnight at 37 °C, then inoculated into tryptic soy broth for 18-20h overnight growth at 37 °C with shaking at 225 rpm. Overnight suspensions were diluted to an OD_595_ of ∼0.05 and placed within the inner 60 wells of a 96-well plate equipped with a lid. To assess growth, isolates were then grown in tryptic soy broth at 37 °C and reads in the GFP channel (excitation: 485/20, emission: 516/20) were taken at 10-min intervals in a BioTek Synergy 2 microplate reader set to moderate continuous shaking for 24 hours. Relative fluorescence units (RFU) of the *P. aeruginosa* strains in tryptic soy broth over 24 hours was plotted using the Growthcurver package in R (*23*).

### Invasion assays of GFP-tagged *P. aeruginosa* strains into the respiratory microbiome strains

An invasion assay in which endpoint measurements of fluorescence in the GFP channel (excitation: 485/20, emission: 516/20) at 24 hours was used to measure the ability of PAO1-GFP and the nine spontaneous antibiotic resistant mutants to invade dense overnight cultures of the six respiratory microbiome strains. The ten *P. aeruginosa* strains (PAO1-GFP, cipR1, cipR2, cipR3, cefR1, cefR2, cefR3, merR1, merR2 and merR3) and six respiratory microbiome strains (*S. epidermidis, R. mucilaginosa, S. gordonii, S. lugdunensis, S. oralis, and S. salivarius*) were grown from glycerol stock on LB Miller agar plates overnight at 37 °C. Single colonies of each *P. aeruginosa* strain were inoculated into the inner 60 wells of a 96-well plate with wells filled with 200uL tryptic soy broth for overnight growth at 37 °C and shaking at 225rpm, for six biological replicates per strain. Single colonies of each respiratory microbiome strain were inoculated into 14 mL falcon tubes filled with 7 mL tryptic soy broth for overnight growth at 37 °C with shaking at 225 rpm, for six biological replicates per strain.

The next day, a black-sided 96-well plate was labelled for each respiratory microbiome strain, and the following plate layout was used to arrange respiratory microbe invasions (Supplementary Figure 1). In brief, each plate contained six replicates of the ten *P. aeruginosa* strains inoculated into six replicates of the respiratory microbiome strain, with the invasion set up as a 1/20 dilution. Three 96-well plates were used for tryptic soy broth controls, in which the same six replicates of the ten *P. aeruginosa* strains were inoculated (1/20 dilution) into sterile tryptic soy broth for three technical replicates of *P. aeruginosa* growth. This paired growth of *P. aeruginosa* invasion inoculations was done in order to calculate *P. aeruginosa* invasion into respiratory microbe culture as a percentage of the growth achieved in rich media in the absence of that respiratory microbe. The plates were incubated for 24 hours at 37 °C with shaking at 225 rpm. After 24 hours, OD_595_ and GFP (excitation: 485/20, emission: 516/20) of the respiratory microbiome invasions and the tryptic soy broth paired growth assays were read in a BioTek Synergy 2 microplate reader.

Plates in which PAO1-GFP did not grow in 24 hours were then incubated for a further 12 hours, and OD_595_ and GFP were measured again at 36 hours. Invasion ability is given as percentage Relative Fluorescence Units (RFU, arbitrary unit) standardised against growth in the absence of the respiratory microbe. Prior to these invasion assays in which endpoint measurements were taken, invasion was first continually monitored in a BioTek Synergy 2 microplate reader taking reads at 10-minute intervals for 24 hours. This preliminary data was used to inform the endpoint assays and produce invasion growth curves for illustrative purposes, which was done using the Growthcurver package in R (*23*).

### Genome sequencing and variant detection of spontaneous antibiotic resistant mutants of *P. aeruginosa*

The ten *P. aeruginosa* strains (PAO1-GFP, cipR1, cipR2, cipR3, cefR1, cefR2, cefR3, merR1, merR2 and merR3) were sequenced using Illumina short-read sequencing. DNA extraction, library preparation and sequencing were performed by MicrobesNG (Birmingham, UK) according to their protocols. Briefly, ∼4-6×10^9^ cells were concentrated and suspended in 500 μl of DNA/RNA Shield (Zymo Research) in 2 mL screw cap tubes, and stored at room temperature prior to sample submission. Between 5-40 μl of cell suspension was lysed with 120 μl TE buffer containing lysozyme (final concentration 0.1 mg/mL) and RNase A at 37 °C for 25 min. Protein digestion was performed using Proteinase K and SDS with incubation at 65 °C for 5 min. Genomic DNA was purified using an equal volume of paramagnetic beads (SPRI beads, Beckman Coulter) and resuspended in EB buffer (10mM Tris-Hcl, pH 8.0). DNA was quantified with QuantiT dsDNA HS kit (ThermoFisher Scientific) using an Eppendorf AF2200 plate reader and diluted to an appropriate concentration for library preparation.

Library preparation was performed using the Nextera XT Library Prep Kit according to the manufacturer’s protocol with modifications (i.e. two-fold increase in input DNA, PCR elongation increased to 45 s). DNA quantification and library preparation were performed using a Hamilton Microlab STAR liquid handling system. Libraries were sequenced using the Illumina NovaSeq6000 platform using a 250 bp paired-end protocol. Read adapter trimming was performed using Trimmomatic version 0.30 (*24*) with a sliding window quality cut-off of Q15. Trimmed reads were aligned to the PAO1-UW reference genome (NCBI RefSeq accession NC_002516.2) (*25*). Variants in PAO1-GFP and the resistant strains were called using the breseq 0.36.1 pipeline (*26*). Variants reported in the main text are those found in the resistant strains only and called at 100% (Supplementary Table 2).

### Spent media assays with Staphylococcus lugdunensis

To better characterise the interaction between *P. aeruginosa* and *S. lugdunensis*, we repeated the invasion assays using clinical isolates of *P. aerugunisa* from two patients where meropenem resistance evolved during infection and the *de novo* resistance mutations are known (Supplementary Table 1) (*27, 28*). A spent media assay was used for this invasion assay as the clinical isolates were not GFP-tagged (*29*). The invasion assay method above was followed with the following alterations. After overnight growth of *S. lugdunensis* cultures, spent media was prepared via centrifugation at 16,000 g/min for 1 minute to pellet the *S. lugdunensis* cells followed by filter sterilisation of the supernatant through a 0.22 μM filter. *P. aeruginosa* strains were inoculated into this cell-free spent media (1/20 dilution) in a 96-well plate as described above alongside paired replicate inoculations into sterile tryptic soy broth. For each clinical *P. aeruginosa* genotype (ST782-WT, *oprD mexR* ST782, ST17-WT, and *oprD* ST17), three biological replicates of three genetically identical isolates from the patient were measured (Supplementary Table 1), and PAO1-GFP was included in these assays as a control.

### Statistical analysis

Statistical analysis was done in R Studio Version 1.1.463 (*30*). ANOVA followed by Dunnett’s test was used to test for differences in invasion ability between the *P. aeruginosa* antibiotic resistant strains and PAO1-GFP. The *DescTools R* package was used to implement the Dunnett’s test (*31*) and significance was determined at p < 0.05. One tailed unpaired t-tests were used to test for differences in growth in the *S. lugdunensis* spent media assays and significance was determined at p < 0.05.

## Results

### Characterisation of *P. aeruginosa* resistant mutants

Spontaneous resistant mutants of GFP-tagged *P. aeruginosa* (PAO1-GFP) were selected at the clinical breakpoints for ciprofloxacin (cipR1, cipR2, cipR3), ceftazidime (cefR1, cefR2, cefR3) and meropenem (merR1, merR2, merR3) (Figure 1) (*21*). Resistance phenotyping, genome sequencing, and growth rate assays were carried out to characterize these resistant mutants (Table 1) (Supplementary Table 2). Selection on ciprofloxacin was associated with the acquisition of mutations in *nfxB*, which encodes NfxB, a transcriptional repressor of the *mexCD*-*oprJ* operon (*32*). Mutations in *nfxB* can upregulate the expression of the multidrug efflux pump MexCD-OprJ (*33, 34*). Efflux pump overexpression is typically associated with a multidrug resistance phenotype, but the *nfxB* mutations were only associated with increases in resistance to ciprofloxacin (Table 1). One ciprofloxacin resistant mutant (cipR2) had a deletion that included both *nfxB* and *morA*, predicted to encode a motility regulator involved in flagellar development and biofilm formation (*35*).

**Figure 1.**
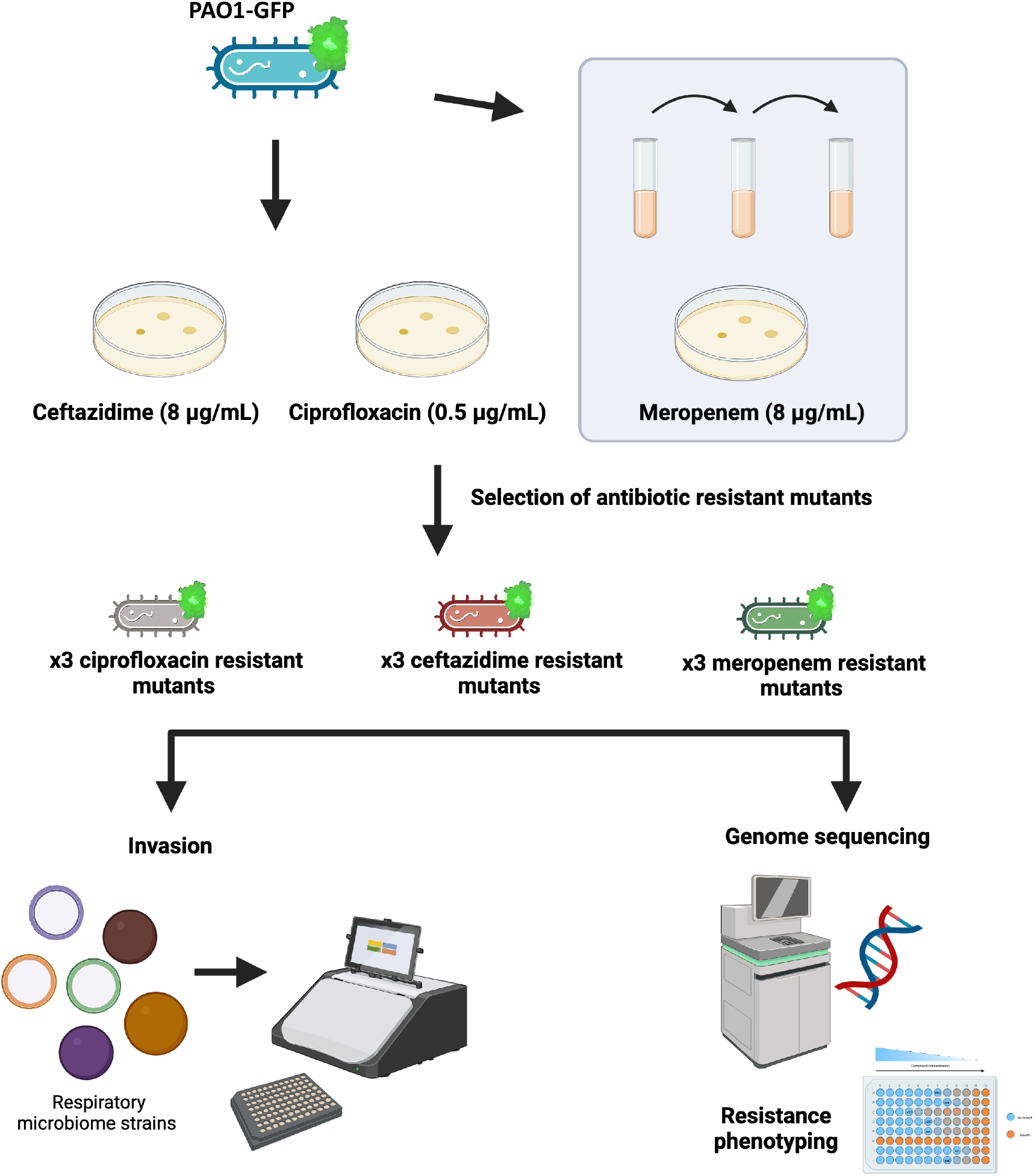
Schematic for selection of *P. aeruginosa* antibiotic resistant mutants, subsequent strain characterization (via genome sequencing and resistance phenotyping), and use of strains in invasion assays.

**Table 1.** Characterisation of PAO1-GFP and resistant mutants (cipR1, cipR2, cipR3, cefR1, cefR2, cefR3, merR1, merR2, merR3).

| Strain | Ciprofloxacin<br>MIC (µg/mL) | Ceftazidime<br>MIC<br>(µg/mL) | Meropenem<br>MIC<br>(µg/mL) | Mutated gene(s) | Mutation type |
| --- | --- | --- | --- | --- | --- |
| PAO1-GFP | 0.25 | 0.5 | 0.5 |  |  |
| cipR1 | 4.0 | 0.5 | 0.5 | *nfxB* | Non-synonymous<br>mutation (R42H) |
| cipR2 | 4.0 | 0.5 | 0.5 | *nfxB*–[*morA*] | Δ4649 bp deletion |
| cipR3 | 2.0 | 0.5 | 0.5 | *nfxB* | Non-synonymous<br>mutation (A33P) |
| cefR1 | 0.25 | 8.0 | 0.5 | *dacB* | Δ5 bp deletion |
| cefR2 | 0.25 | 4.0 | 0.5 | [*PA3046*]–[*dacB*] | Δ1249 bp deletion |
| cefR3 | 0.25 | 8.0 | 0.5 | *dacB* | Non-synonymous<br>mutation (A343E) |
| merR1 | 0.5 | 2.0 | 8.0 | [*phoQ*]–<br>[*PA1181*]–<br>[*PA1182*] | Δ4122 bp deletion |
|  |  |  |  | *nalD* | Non-synonymous<br>mutation (T11N) |
| merR2 | 1 | 2.0 | 8.0 | *mexR* | Δ12 bp deletion |
|  |  |  |  | *phoQ* | Non-synonymous<br>mutation (V382G) |
| merR3 | 0.5 | 1.0 | 8.0 | *oprD* | Δ11 bp deletion |
|  |  |  |  | *phoQ* | Non-synonymous<br>mutation (V260G) |
|  |  |  |  | *nalD* | Non-synonymous<br>mutation (T11P) |

Selection on ceftazidime was associated with the acquisition of mutations in *dacB* that are known to cause the overexpression of the AmpC β-lactamase and increased resistance to ceftazidime (*36*). The meropenem resistant mutants (merR1, merR2, merR3) were selected through a short daily passage experiment rather than from overnight culture plating and had a larger number of variants than the ciprofloxacin and ceftazidime resistant mutants (Figure 1). The genetic basis of meropenem resistance in *P. aeruginosa* is complex and clinical resistance levels require two or more mutations (*37, 38*), this is reflected in the increased complexity of the meropenem resistant mutants that we isolated. Meropenem resistant mutants carried mutations in genes such as *phoQ*, *mexR* and *nalD* that are known to be associated with elevated expression of multidrug efflux pumps (*39, 40*). One of the mutants (merR3) also carried a deletion in *oprD,* which encodes the outer porin OprD that can serve as a channel to let meropenem into the cell (*37, 41*). These mutations in genes involved in multidrug efflux pump regulation likely explain why meropenem resistance was associated with increased resistance to both ceftazidime and ciprofloxacin too (Table 1). The ciprofloxacin and meropenem resistant mutants demonstrated decreased growth compared to PAO1-GFP in rich media (as measured by RFU over time), whereas growth of strains with ceftazidime resistance was more comparable to PAO1-GFP (Figure 2).

**Figure 2.**
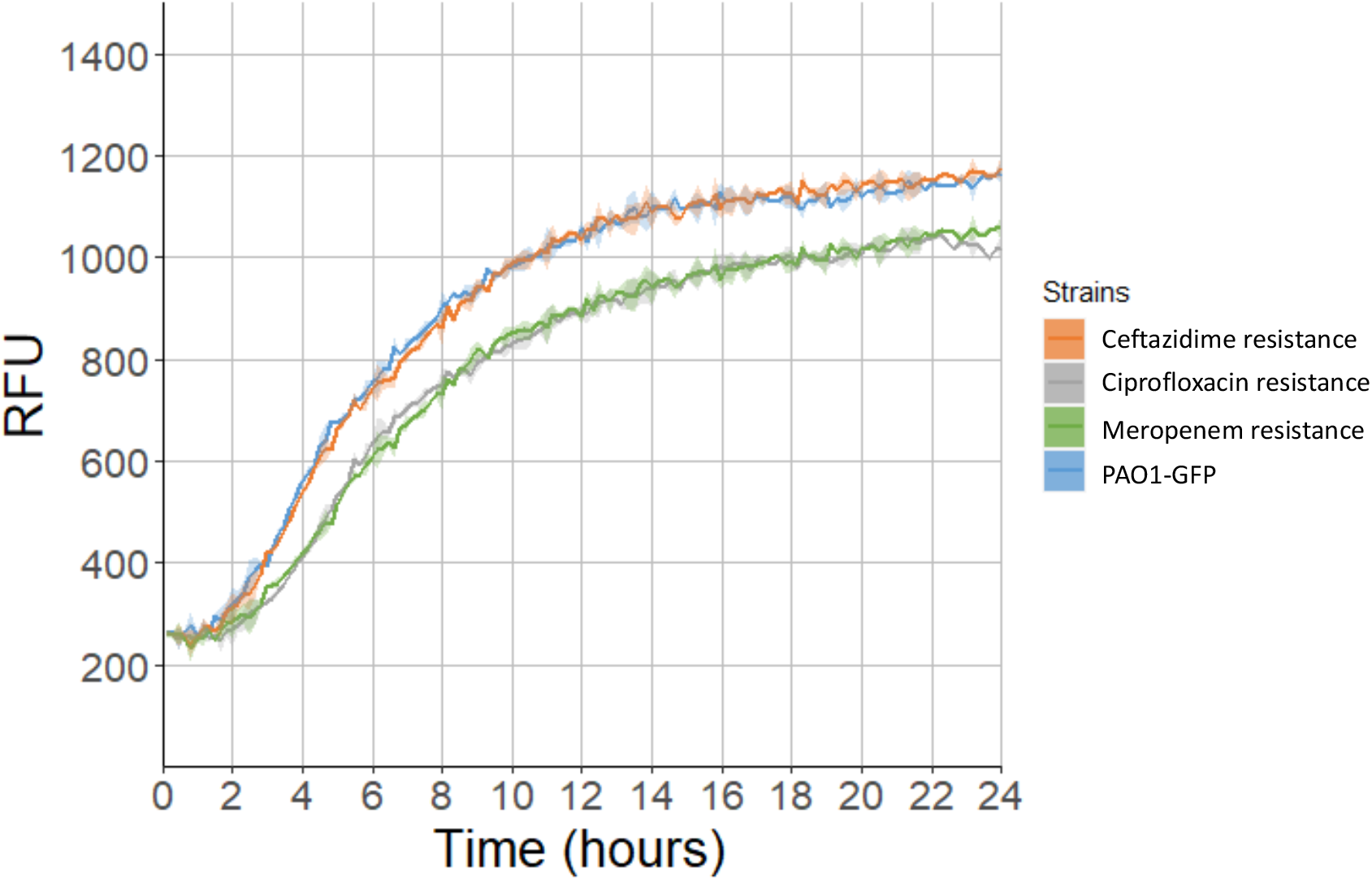
Growth rate assays of the ceftazidime (orange), ciprofloxacin (grey) and meropenem (green) resistant strains compared to PAO1-GFP (blue) in tryptic soy broth in the absence of respiratory microbes. Growth was measured as relative fluorescence units (RFU) in the GFP channel over a period of 24 hours, and data was plotted using the Growthcurver package in R (*23*).

### Invasion assays of *P. aeruginosa* resistant mutants into the respiratory microbiome

We measured the ability of the *P. aeruginosa* resistant mutants to invade the pre-established respiratory microbiome strains by measuring GFP at 24 hours after *P. aeruginosa* inoculation (Figure 3). These invasion assays were standardised against paired growth of *P. aeruginosa* in tryptic soy broth (in absence of a respiratory microbe), and invasion assay results are shown as percentage growth in respiratory microbe invasion compared to the media control.

**Figure 3.**
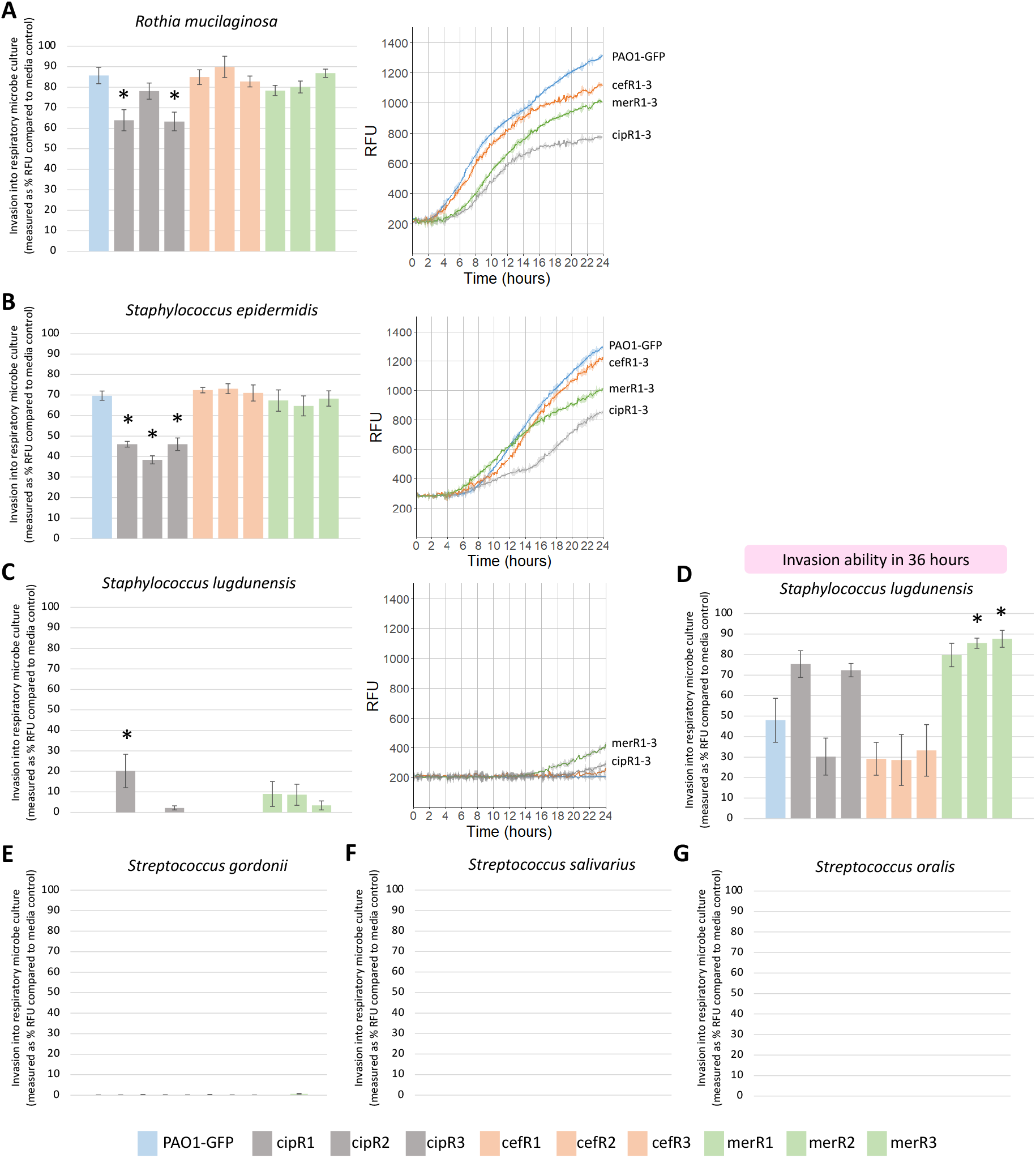
Invasion of *P. aeruginosa* into respiratory microbiome strains in 24 hours (A, B, C, E, F, G) and 36 hours (D). Each bar chart shows invasion ability measured as percentage RFU compared to the media control at that same timepoint, for invasion into pre-established cultures of: (A) *R. mucilaginosa*, (B) *S. epidermidis*, (C) *S. lugdunensis* (24 hours), (D) *S. lugdunensis* (36 hours), (E) *S. gordonii*, (F) *S. salivarius*, and (G) *S. oralis*. Values obtained in *S. gordonii*, *S. salivarius*, and *S. oralis* were considered negligible and below the limits of the assay (below 2%). ANOVA followed by Dunnett’s test was carried out to compare invasion ability of PAO1-GFP to each resistant mutant, and significance as p < 0.05 is indicated by an asterix (*). Bar charts show the mean of six biological replicates +/− standard error of the mean. For the invasion assays in which *P. aeruginosa* was able to successfully invade (*R. mucilaginosa, S. epidermidis, S. lugdunensis*), pilot growth curves are shown to the right of the bar chart for illustrative purposes (PAO1-GFP = blue, cipR1-R3 = gray, cefR1-R3 = orange, merR1-R3 = green). These were plotted using the Growthcurver package in R (*23*).

All respiratory microbe cultures provided some degree of inhibition to *P. aeruginosa* growth, and this effect was particularly strong for the *Streptococcus* species and *S. lugdunensis* (Figure 3). *P. aeruginosa* was best able to invade *R. mucilaginosa* (Figure 3A), followed by *S. epidermidis* (Figure 3B). Ciprofloxacin resistant mutants had impaired invasion ability into both *R. mucilaginosa* and *S. epidermidis* (Figure 3A, 2B), suggesting a clear cost to resistance in this context. On the other hand, the resistant mutants demonstrated an increased invasion ability into cultures of *S. lugdunensis* (Figure 3C). *P. aeruginosa* (PAO1-GFP) was unable to invade *S. lugdunensis* in 24 hours, but five of the resistant mutants (cipR1, cipR3, merR1, merR2, merR3) were able to grow in this same timeframe (defined as at least 2% of RFU in media in absence of *S. lugdunensis*) (Figure 3C).

The invasion of the resistant mutants into *S. lugdunensis* was near our lower limit of detection (Figure 3C), and so we extended this assay to 36 hours. In 36 hours, PAO1-GFP was now able to invade *S. lugdunensis* (Figure 2D). The five resistant mutants (cipR1, cipR3, merR1, merR2, merR3) demonstrated a trend towards increased invasion ability (Figure 3D), suggesting a benefit to resistance in this context. It should be noted that statistical significance (p < 0.05) as assessed via the Dunnett’s test was only observed for cipR1 at 24 hours and merR1/merR2 at 36 hours (Figure 3D). Finally, the *Streptococccus* spp. (*S. gordonii, S. salivarius, S. oralis*) showed strong inhibition of *P. aeruginosa* growth, which was maintained regardless of the resistance genotype (Figure 3E, F, G). *P. aeruginosa* was unable to invade in 24 hours, and this effect was preserved when we extended the assay to 36 hours (Supplementary Figure 2). The three ceftazidime resistant mutants (cefR1, cefR2, cefR3) associated with mutations in *dacB* were not found to alter invasion into any of the six respiratory microbes.

### Invasion of clinical isolates of P. aeruginosa into Staphylococcus lugdunensis

Invasion of *P. aeruginosa* into *S. lugdunensis* was an interesting example of where antibiotic resistance seemed to improve the ability of *P. aeruginosa* to invade (Figure 3C, 2D). To better characterise this interaction, we repeated these invasion assays using isolates from two patients where meropenem resistance evolved during infection due to the acquisition of *de novo* mutations (Supplementary Table 1) (*27, 28*). These isolates were obtained from two hospitalised patients that were previously recruited as part of an observational trial into resistance evolution during *P. aeruginosa* infections in intensive care (*42, 43*), and are described in detail here: (*27, 28*). In brief, one patient was colonised by Sequence Type (ST) 782 *P. aeruginosa* (ST782-WT), and increased resistance to meropenem evolved via mutation to *oprD* and *mexR* (*oprD mexR* ST782) (*28*). The second patient was colonised by ST17 *P. aeruginosa* (ST17-WT), and increased resistance to meropenem evolved via mutation to *oprD* (*oprD* ST17) (*27*). These assays were carried out as spent media assays, in which *P. aeruginosa* invasion ability was measured using OD_595_ reads, as these clinical isolates were not GFP-tagged (Figure 4A). PAO1-GFP was tested in parallel in this assay.

**Figure 4.**
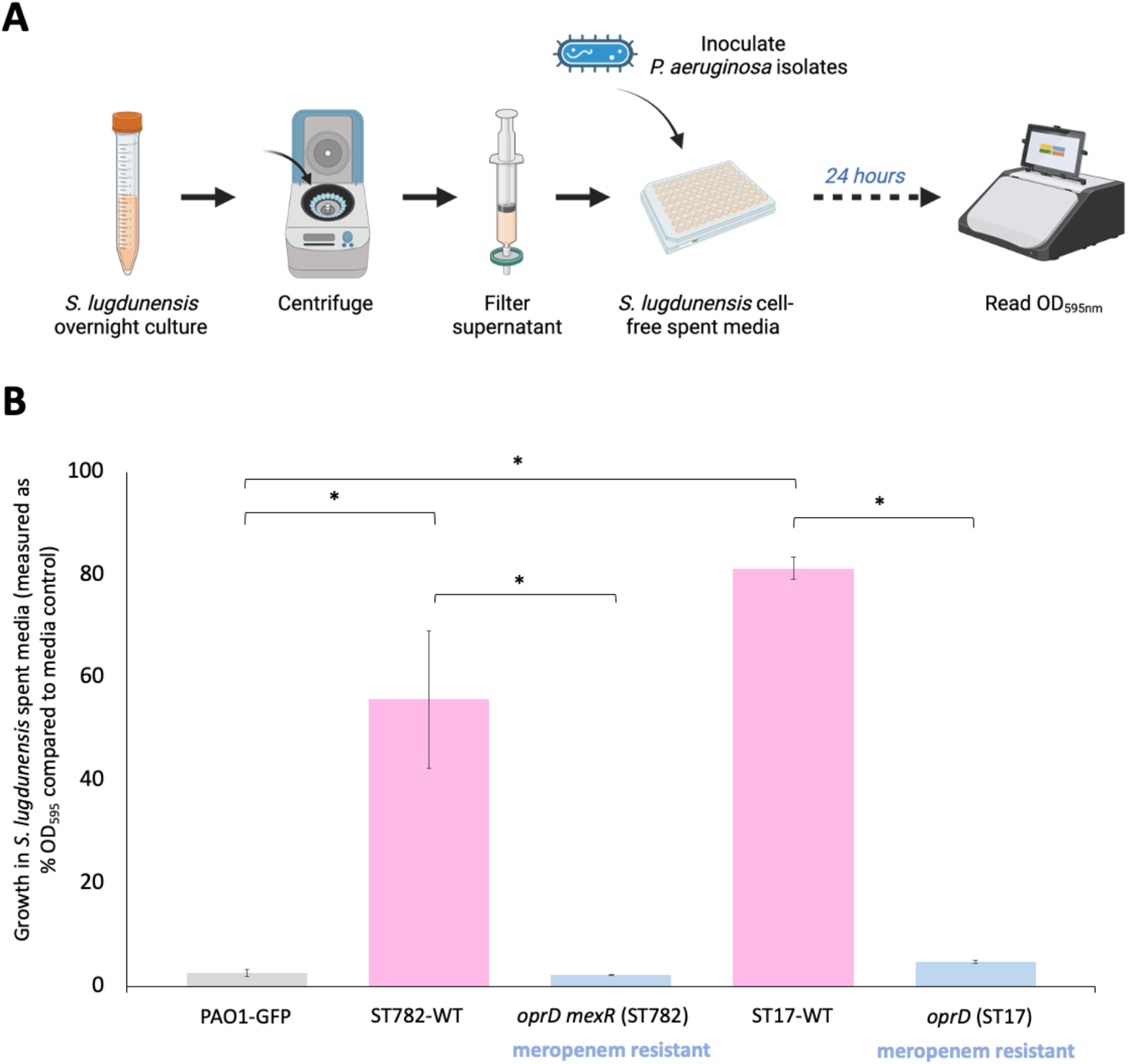
(A) Schematic of *S. lugdunensis* spent media assay. (B) Growth of PAO1-GFP and clinical *P. aeruginosa* isolates in *S. lugdunensis* spent media, measured as percentage OD_595_ in *S. lugdunensis* spent media compared to the media control at 24 hours. For each clinical *P. aeruginosa* genotype, three biological replicates of three isolates were measured and the values are plotted as mean +/− standard error of the mean. For PAO1-GFP, three biological replicates were measured. One tailed unpaired t-tests were carried out to compare growth in *S. lugdunensis* spent media between PAO1-GFP and the two clinical WT strains (ST782-WT, ST17-WT) and to compare between each resistant mutant to its WT background. Significance as p < 0.05 is indicated by an asterix (*).

*S. lugdunensis* spent media was able to inhibit PAO1-GFP growth over 24 hours (Figure 4B), a similar effect as observed in *S. lugdunensis* culture (Figure 3C), suggesting the inhibition mediated by *S. lugdunensis* is independent of *P. aeruginosa* detection or contact. However, this inhibitory effect was weak for the clinical isolates ST17-WT and ST782-WT, which were able to grow to ∼80% and ∼55% of the OD_595nm_ reached in the media control respectively (Figure 4B). Conversely, mutations that increased meropenem resistance during infection were associated with a stronger inhibitory effect for growth in *S. lugdunensis* spent media. The ST17 isolates with mutations in *oprD* showed dramatically reduced growth in *S. lugdunensis* spent media compared to ST17-WT, as did ST782 isolates with mutations in both *oprD* and *mexR* (Figure 4B). This suggests that mutation to *oprD* in merR3 did not likely contribute to the improved invasion ability of meropenem resistant PAO1-GFP mutants (Figure 3).

## Discussion

The emergence and spread of antibiotic resistance in bacterial pathogens is responsible for a range of problems on a global scale (*1, 2*). It is key that we understand the ecological consequences of antibiotic resistance, so that we can predict and appropriately respond to these problems. Here we show that antibiotic resistance alters the ability of *P. aeruginosa* to invade the respiratory microbiome, in both positive and negative directions, which are *P. aeruginosa* genotype and respiratory microbe identity dependant. We hypothesised at the onset of this work that although antibiotic resistance is often associated with a fitness cost (*5–7*), there may be members of the respiratory microbiome whereby resistance provides *P. aeruginosa* with an enhanced invasion ability into. Indeed, we found that *P. aeruginosa* resistant mutants were able to invade *S. lugdunensis* in a period of 24 hours, whereas the wildtype could not (Figure 3C).

The two ciprofloxacin resistant mutants (cipR1, cipR3) better able to invade *S. lugdunensis* both had mutations in *nfxB* (Table 1). Mutations in *nfxB* can upregulate the expression of the multidrug efflux pump MexCD-OprJ (*33, 34*), suggesting overexpression of MexCD-OprJ may be beneficial for invasion into *S. lugdunensis.* Interestingly, the third ciprofloxacin resistant mutant (cipR2) that had a deletion spanning *nfxB* and *morA* did not demonstrate an improved invasion ability (Figure 3C, 3D). This suggests the phenotypes regulated by MorA, which include flagella development and biofilm formation (*35*), may be important for invasion into *S. lugdunensis*, and the fitness costs associated with this deletion may be enhanced under these conditions. The three meropenem resistant mutants (merR1, merR2, merR3) better able to invade *S. lugdunensis* similarly had mutations in genes involved in multidrug efflux pump regulation (Table 1). Both *nalD* and *mexR* encode repressors of the multidrug efflux pump MexAB-OprM, suggesting overexpression of MexAB-OprM is also beneficial for invasion into *S. lugdunensis.* Similarly, the PhoP-PhoQ two-component regulatory system has been identified with a role in regulating multidrug efflux pumps, among a wide variety of other regulatory activities (*44–48*), and the meropenem resistant mutants also had mutations in *phoQ* (Table 1).

That increased expression of efflux pumps (e.g. MexCD-OprJ and MexAB-OprM) may help facilitate invasion into *S. lugdunensis* led us to hypothesise that *S. lugdunensis* is secreting an inhibitory molecule that can be exported by these pumps. In support of this hypothesis, PAO1-GFP was also unable to grow in the cell-free spent media of *S. lugdunensis* (Figure 4B). Lugdunin has been characterised as a novel thiazolidine-containing cyclic peptide antibiotic that can be produced by *S. lugdunensis* strains. Lugdunin has not been shown to be significantly active against *P. aeruginosa* (*49, 50*), suggesting another secreted molecule may be responsible for this inhibitory phenotype. While the enhanced invasion phenotype of the efflux pump regulatory mutants fits well with this hypothesis, it is also possible that some of the *S. lugdunensis* inhibition could be mediated by depletion of metabolites required by *P. aeruginosa*. Under this model, the growth of *S. lugdunensis* restricts *P. aeruginosa* indirectly via the consumption of nutrients needed for *P. aeruginosa* growth in this environment. Interestingly, recent work has demonstrated how the ability of multi-drug resistant *Escherichia coli* lineages to displace commensal *E. coli* in the gut microbiome can be facilitated by encoding a high genetic diversity of carbohydrate metabolism genes, which can help provide a competitive advantage to invade in these environments (*51*).

A deletion in *oprD* was found in one of the meropenem resistant mutants (merR3) (Table 1)*, oprD* encodes a well-characterised channel for meropenem influx. We were able to further explore this using clinical isolates that evolved high level meropenem resistance in response to treatment during infection via mutation to *oprD* (Figure 4) (*27, 28*). Compared to PAO1-GFP, the clinical isolates (*P. aeruginosa* ST17 and ST782) were able to grow significantly better in the cell-free spent media of *S. lugdunensis* (Figure 4B). However, ST17 and ST782 isolates with *oprD* mutation showed dramatically reduced invasion ability, including the ST782 isolates that also possessed mutation to *mexR* (Figure 4B). In *P. aeruginosa*, *oprD* mutants have shown varied results with regards to a fitness cost. Some studies have shown no costs, and even enhanced survival in infected mouse models (*52*), while other studies have shown meropenem resistance is costly under some conditions, but not others (*27, 28*). Here we see that *oprD* mutations may be associated with the cost of reduced fitness in the presence of *S. lugdunensis* (*53*). Given that OprD is a porin, we speculate that loss of OprD may hinder the ability of clinical isolates to acquire metabolites that are effectively depleted by *S. lugdunensis.* This speculation lends support to a role of nutrients in helping mediate *P. aeruginosa* inhibition by *S. lugdunensis*. These results also suggest that mutation to *oprD* did not likely contribute to the improved invasion ability of merR3, and further to this, suggests that it may be harder for *oprD* mutants to transmit to different patients that have *S. lugdunensis* in their respiratory microbiome.

*P. aeruginosa* was best able to invade *R. mucilaginosa* and *S. epidermidis* cultures, with PAO1-GFP reaching ∼86% ± 4% and ∼70% ± 2% of the RFU reached in the media controls respectively (Figure 3A, 3B). *P. aeruginosa* can utilize *R. mucilaginosa*-produced metabolites as precursors for the generation of primary metabolites (*54*), and that *R. mucilagnosa* presence can help facilitate *P. aeruginosa* growth via cross-feeding may help explain the high growth of *P. aeruginosa* in *R. mucilaginosa* culture invasion. We found that the ciprofloxacin resistant *nfxB* mutants had impaired invasion ability into *R. mucilaginosa* (Figure 3A), demonstrating a clear cost to antibiotic resistance in this scenario. We hypothesise that the reduced invasion ability of *nfxB* mutants in the presence of *R. mucilaginosa* may be due to increased efflux of the *R. mucilaginosa*-produced metabolites that support *P. aeruginosa* growth via the MexCD-OprJ efflux pump. The ciprofloxacin resistant mutants also showed impaired invasion ability in *S. epidermidis* (Figure 3B), leading us to speculate a possible role of MexCD-OprJ overexpression in the efflux of beneficial *S. epidermidis*-produced metabolites or that pump overexpression may impair the ability of *P. aeruginosa* to acquire nutrients depleted by *S. epidermidis*.

Efflux pumps are involved in a variety of biological roles (*55, 56*), and can extrude a wide range of substrates in addition to antibiotics, including metabolites, organic solvents, and cell signalling molecules (*57–59*). Overexpression of the MexCD-OprJ efflux pump has been linked with multiple effects, including impaired quorum sensing (*60*), reductions in type III secretion (*61*), and broad-scale proteome changes (*62*). Similarly, the MexAB-OprM efflux pump has been linked with several roles, including in quorum sensing (*63, 64*) and siderophore secretion (*65*). The ciprofloxacin resistant *nfxB* mutants displayed a particularly interesting phenotype, with reduced invasion ability in *R. mucilaginosa* and improved invasion ability in *S. lugdunensis*. One mechanism that could explain this is that MexCD-OprJ is able to efflux the *R. mucilaginosa*-produced metabolites that support *P. aeruginosa* growth, as well as an *S. lugdunensis-*produced inhibitory molecule that restricts *P. aeruginosa* growth (Figure 5). This could account for the interesting observed phenotype where *nfxB* mutants have increased resistance to ciprofloxacin and an improved invasion ability in *S. lugdunensis,* but demonstrate a clear cost to *nfxB* mutation in the presence of *R. mucilaginosa* (Figure 5). Future work would need to validate this mechanism, particularly considering the other varied roles that MexCD-OprJ overexpression can play, and distinguishing between inhibition mediated by presence of a secreted inhibitory molecule versus via absence of a substrate needed to facilitate growth.

**Figure 5.**
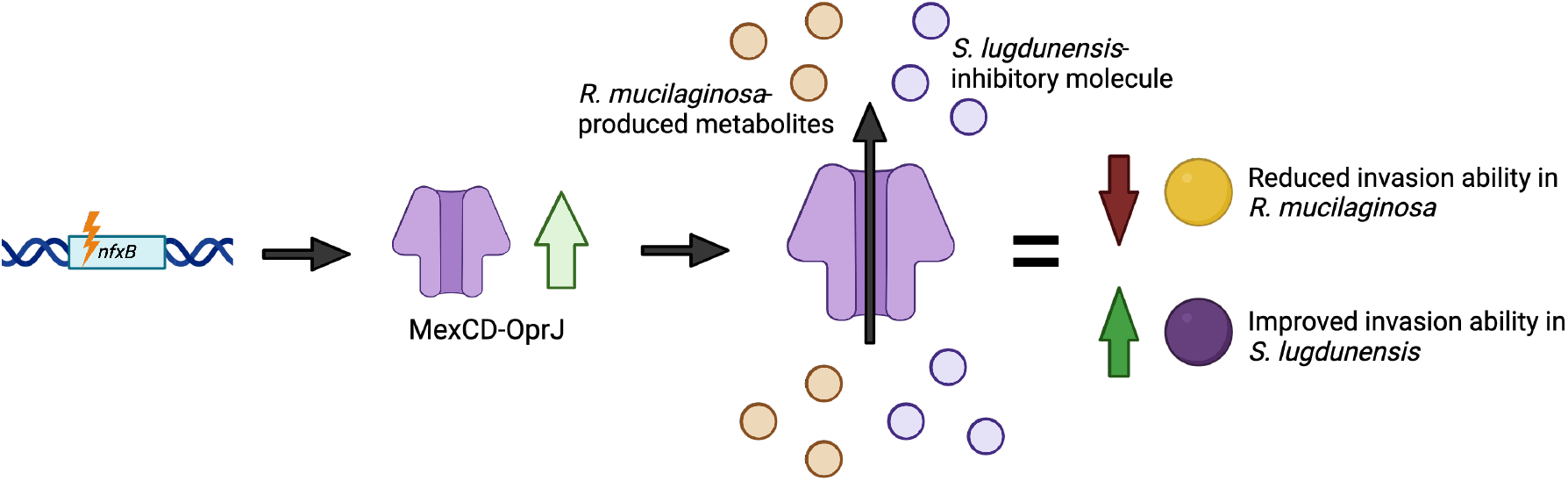
Hypothetical mechanism by which ciprofloxacin resistant *nfxB* mutants could demonstrate reduced invasion ability in *R. mucilaginosa* and improved invasion ability in *S. lugdunensis*.

Finally, we identified three *Streptococcus* species (*S. gordonii, S. salivarius, S. oralis*) that *P. aeruginosa* was unable to invade, regardless of resistance genotype (Figure 3E, 3F, 3G). These streptococcal species are known for their ability to inhibit *P. aeruginosa* through a variety of mechanisms (*66–68*), including media acidification (*69*), hydrogen peroxide production (*70*), and through production of a variety of bacteriocin-like inhibitory substances (*71*). *S. salivarius* strains have been developed for oral probiotic applications that are commercially available (*72*). Our research adds to this body of literature and demonstrates that successful inhibition of *P. aeruginosa* still occurs for a variety of resistance genotypes that have been generated by selection on important anti-pseudomonal drugs.

## Conclusions

In summary, we find that commensal respiratory microbe strains tend to inhibit the growth of *P. aeruginosa*, and antibiotic resistance is a double-edged sword that can either help or hinder the ability of *P. aeruginosa* to overcome this inhibition. Antibiotic resistance facilitates the invasion of *P. aeruginosa* into *S. lugdunensis,* yet impairs invasion into *R. mucilaginosa* and *S. epidermidis*. *Streptococcus* species provide the strongest inhibition to *P. aeruginosa* invasion, and this is maintained regardless of antibiotic resistance genotype. The ciprofloxacin resistant *P. aeruginosa* strains associated with mutations in *nfxB* showed altered invasion into *S. lugdunensis, S. epidermidis* and *R. mucilaginosa*, and this altered invasion ability was in both directions depending on respiratory microbe identity. On the other hand, the ceftazidime resistant strains associated with mutations in *dacB* were not found to alter invasion into any of the six respiratory microbes. Looking to the future, we argue that understanding this interplay between antibiotic resistance and interactions in the microbiome has clear significance for understanding the potential of resistant strains to transmit between patients. Attempts to manipulate the respiratory microbiome should focus on promoting the growth of commensals that can provide robust inhibition of both wildtype and resistant mutant strains. We hope that building this knowledge will help us identify high-risk *P. aeruginosa* resistant mutants for patient transmission, and further to this, help us identify microbiome signatures that may be more susceptible to antibiotic-resistant pathogen invasion.

## Acknowledgements

R.M.W was supported by the George Grosvenor Freeman Fellowship by Examination in Sciences, Magdalen College (Oxford), and by the Calleva Research Centre for Evolution and Human Sciences at Magdalen College, Oxford. S.S. was supported by the Calleva Research Centre for Evolution and Human Sciences at Magdalen College, Oxford. This work was supported by the Department of Biology (University of Oxford) financial support for MBiol projects, received by S.L. This work was additionally supported by grants to DRG (Academy of Medical Sciences Springboard Award: SBF007\100096, BBSRC: BB/X007979/1). We would like to thank Jason L Brown and Gordon Ramage (Oral Sciences Research Group, University of Glasgow), Angela Nobbs (University of Bristol), and Josh Thomas and Ashleigh Griffin (University of Oxford) for strains provided and used in this study. We would like to thank MicrobesNG (Birmingham, UK) for the generation and initial processing of sequencing data. We would like to thank Alexandre Figueiredo and Erik Bakkeren for comments on the manuscript. Figures within this publication were created with Biorender.com (Figure 1, Figure 3A, Figure 4).

## Supplementary Information

**Supplementary Table 1.** Strain lists.

**Supplementary Table 2.** Variants found in resistant strains compared to the PAO1-GFP background. DEL; deletion, SNP; single nucleotide polymorphism.

**Supplementary Figure 1.** Plate layout for invasion assays.

**Supplementary Figure 2.** Invasion of *P. aeruginosa* strains into *Streptococcus* species over 36 hours.

