## Supplementary Information for "Antibiotic resistance alters the ability of *Pseudomonas aeruginosa* to invade the respiratory microbiome"

### **Supplementary Figure 1.** Plate layout for invasion assays.

The respiratory microbe cultures were arranged into a 96-well plate such that six replicate cultures were present in each of the 10 inner well columns of the plate, as is shown below of the example of 'SE':

| | 1 | 2 | 3 | 4 | 5 | 6 | 7 | 8 | 9 | 10 | 11 | 12 |
|---|---|---|---|---|---|---|---|---|---|---|---|---|
| A | | | | | | | | | | | | |
| B | rep 1 | SE-1 | SE-1 | SE-1 | SE-1 | SE-1 | SE-1 | SE-1 | SE-1 | SE-1 | SE-1 | |
| C | rep 2 | SE-2 | SE-2 | SE-2 | SE-2 | SE-2 | SE-2 | SE-2 | SE-2 | SE-2 | SE-2 | |
| D | rep 3 | SE-3 | SE-3 | SE-3 | SE-3 | SE-3 | SE-3 | SE-3 | SE-3 | SE-3 | SE-3 | |
| E | rep 4 | SE-4 | SE-4 | SE-4 | SE-4 | SE-4 | SE-4 | SE-4 | SE-4 | SE-4 | SE-4 | |
| F | rep 5 | SE-5 | SE-5 | SE-5 | SE-5 | SE-5 | SE-5 | SE-5 | SE-5 | SE-5 | SE-5 | |
| G | rep 6 | SE-6 | SE-6 | SE-6 | SE-6 | SE-6 | SE-6 | SE-6 | SE-6 | SE-6 | SE-6 | |
| H | | | | | | | | | | | | |

The *P. aeruginosa* strains were then inoculated into this plate for six replicates of each strain as shown below:

| | 1 | 2 | 3 | 4 | 5 | 6 | 7 | 8 | 9 | 10 | 11 | 12 |
|---|---|---|---|---|---|---|---|---|---|---|---|---|
| A | | | | | | | | | | | | |
| B | rep 1 | PAO1-GFP-1 | cipR1-1 | cipR2-1 | cipR3-1 | cefR1-1 | cefR2-1 | cefR3-1 | merR1-1 | merR2-1 | merR3-1 | |
| C | rep 2 | PAO1-GFP-2 | cipR1-2 | cipR2-2 | cipR3-2 | cefR1-2 | cefR2-2 | cefR3-2 | merR1-2 | merR2-2 | merR3-2 | |
| D | rep 3 | PAO1-GFP-3 | cipR1-3 | cipR2-3 | cipR3-3 | cefR1-3 | cefR2-3 | cefR3-3 | merR1-3 | merR2-3 | merR3-3 | |
| E | rep 4 | PAO1-GFP-4 | cipR1-4 | cipR2-4 | cipR3-4 | cefR1-4 | cefR2-4 | cefR3-4 | merR1-4 | merR2-4 | merR3-4 | |
| F | rep 5 | PAO1-GFP-5 | cipR1-5 | cipR2-5 | cipR3-5 | cefR1-5 | cefR2-5 | cefR3-5 | merR1-5 | merR2-5 | merR3-5 | |
| G | rep 6 | PAO1-GFP-6 | cipR1-6 | cipR2-6 | cipR3-6 | cefR1-6 | cefR2-6 | cefR3-6 | merR1-6 | merR2-6 | merR3-6 | |
| H | | | | | | | | | | | | |

Key

Outer wells filled with media

Inner wells used for invasion assay

**Supplementary Figure 2.** Invasion of *P. aeruginosa* strains into *Streptococcus* species over 36 hours.

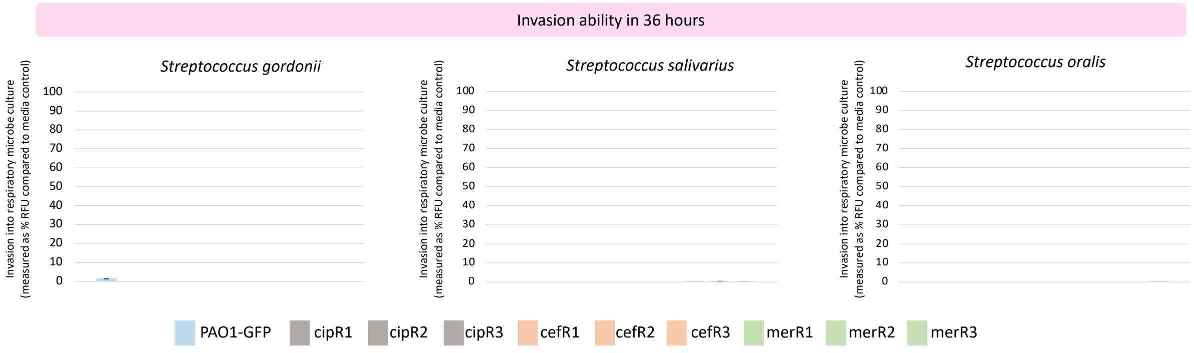
